# Injury-induced Erk1/2 signaling enhances Ca^2+^ activity and is necessary for regeneration of spinal cord and skeletal muscle

**DOI:** 10.1101/2021.09.12.459948

**Authors:** Jacqueline B. Levin, Laura N. Borodinsky

## Abstract

The transition of stem cells from quiescence to proliferation enables tissues to self-repair. The signaling mechanisms driving these stem-cell-status decisions are still unclear. Ca^2+^ and the extracellular signal-regulated kinase (Erk1/2) are two signaling pathways that have the potential to coordinate multiple signals to promote a specific cellular response. They both play important roles during development but their roles during regeneration are not fully deciphered. Here we show in *Xenopus laevis* larvae that both Ca^2+^ and Erk1/2 signaling pathways are activated after tail amputation. In response to injury, we find that Erk1/2 signaling is activated in neural and muscle stem cells and is necessary for spinal cord and skeletal muscle regeneration. Finally, we show *in vivo* that Erk1/2 action is necessary for an injury-induced increase in intracellular store-dependent Ca^2+^ dynamics in skeletal muscle-associated tissues but that in spinal cord, injury increases Ca^2+^ influx-dependent Ca^2+^ activity independent of Erk1/2 signaling. This study suggests that precise temporal and tissue-specific activation of Ca^2+^ and Erk1/2 pathways is essential for regulating tissue regeneration.

## INTRODUCTION

The ability of organisms to replace lost or damaged tissue is essential to organ homeostasis, and deficits in this ability hinder recovery from injury, disease, or other stressors. Regeneration in humans is limited, varies by tissue type, and generally decreases with age (Chen et al., 2020; Gurtner et al., 2008; Tanaka & Ferretti, 2009; Uygur & Lee, 2016). In humans, most new cells are generated from tissue-specific stem cells that are precisely controlled to allow them to quickly activate to replace injured tissue. Understanding the signaling mechanisms that are engaged in these cells is vital to devising therapeutic strategies to enhance tissue regeneration. Many species, including *Xenopus laevis* larvae, are more proficient at regeneration than humans, providing us opportunities to gain insight into optimizing stem cell function by investigating their intrinsic mechanisms that participate in tissue regeneration.

Both the Ca^2+^ and Ras/Raf/MEK1/2/ERK1/2 signaling pathways are ubiquitous and responsive to numerous extracellular signals, causing a diverse array of downstream effects and making them ideal candidates to integrate multiple stimuli into a discrete cellular response. The canonical ERK1/2 signaling pathway initiates when an extracellular ligand binds a receptor tyrosine kinase, usually a growth factor receptor, leading to activation of Ras which promotes Raf serine/threonine kinase activity and sequential activation of MEK1/2 and ERK1/2 by phosphorylation (English et al., 1999). Phosphatases and other enzymes are able to modify activation at every level of this pathway, leading to signal diversity (Roskoski, 2012). Ca^2+^ dynamics are also spatiotemporally controlled via level of expression and subcellular localization of channels, pumps and buffer proteins that bind to and sequester Ca^2+^ (Carafoli & Krebs, 2016).

Previous studies have implicated both Ca^2+^ and ERK1/2 signaling in regeneration. Altering intracellular Ca^2+^ stores interferes with regeneration of *Xenopus* larval tail (Tu & Borodinsky, 2014) and zebrafish fin (Yoo et al., 2012) through unknown mechanisms. In addition, Fgfr1, which often acts by promoting Erk1/2 signaling, is necessary for regeneration of the *Xenopus* larval tail (Lin & Slack, 2008) and Fgf signaling promotes neural progenitor proliferation and functional recovery after spinal cord injury in zebrafish (Goldshmit et al., 2012) and mice (Goldshmit et al., 2014).

Examining the signaling mechanisms in muscle satellite cells (skeletal muscle stem cells) provides more clues about how ERK1/2 signaling may affect stem cell function. In mouse muscle satellite cells, sustained ERK1/2 activation promoted by SHP2 action (Griger et al., 2017), and the activation of FGF signaling by knocking down its inhibitor, SPRY1 (Chakkalakal et al., 2012; Shea et al., 2010), both promote satellite cell proliferation. Reinstating FGF inhibition is necessary to reverse activation to maintain the satellite cell pool and thereby regenerative capacity (Shea et al., 2010). The receptor tyrosine kinase c-Met, is also expressed in mouse muscle satellite cells (Allen et al., 1995; Cornelison & Wold, 1997), and is necessary for their enhanced activation in response to HGF signaling (Rodgers et al., 2014, 2017). In addition, ERK1/2 activation prevents myogenic differentiation *in vitro* (Kook et al., 2008; Koyama et al., 2008; Milasincic et al., 1996; Wu et al., 2015; Yang et al., 2006), and inhibiting FGF signaling promotes presumptive differentiation of myocytes *in vivo* in chick embryo (Michailovici et al., 2014). These studies suggest that growth factor-induced ERK1/2 signaling may be a molecular switch that controls both the transition between quiescent muscle satellite cells and intermediate progenitors, and the transition between intermediate progenitors and differentiated myocytes. However, the precise role of ERK1/2 and Ca^2+^ dynamics during regeneration and the mechanisms regulating their spatiotemporal profiles are still unclear.

Here we discovered that Erk1/2 signaling is activated in the neural and muscle stem cells of regenerating tissues and is necessary for the regeneration of both spinal cord and muscle. We determined the spatiotemporal pattern of Ca^2+^ activity *in vivo* during regeneration and its interaction with Erk1/2 signaling in a tissue-specific manner.

## RESULTS

### Injury activates Erk1/2 signaling in neural and muscle stem cells and is necessary for cell proliferation in regenerating tissues

Sox2-expressing neural stem cells (Gaete et al., 2012) and Pax7-expressing muscle satellite cells (Chen et al., 2006) are the source of regenerated spinal cord and muscle, respectively, following tail amputation in *Xenopus laevis* larvae. To assess signaling in spinal cord cells following injury, tails of *Xenopus laevis* larvae were processed at 20 min post-amputation for transverse sectioning followed by immunostaining. Our data show that Erk1/2 signaling is acutely activated in both Sox2+ and Sox2-spinal cord cells 50 to 250 µm from the amputation compared to non-amputated sibling larvae (Figure 1). In contrast, at least 300 µm anterior to the amputation, pErk1/2 activation in amputated larvae is similar to non-amputated siblings, showing a modest injury-induced increase only in Sox2-cells between 350 to 400 µm anterior to the injury (Figure 1). The data suggest an injury-induced recruitment of Erk1/2 signaling in spinal cord cells adjacent to the amputation including neural stem cells.

**Figure 1.**
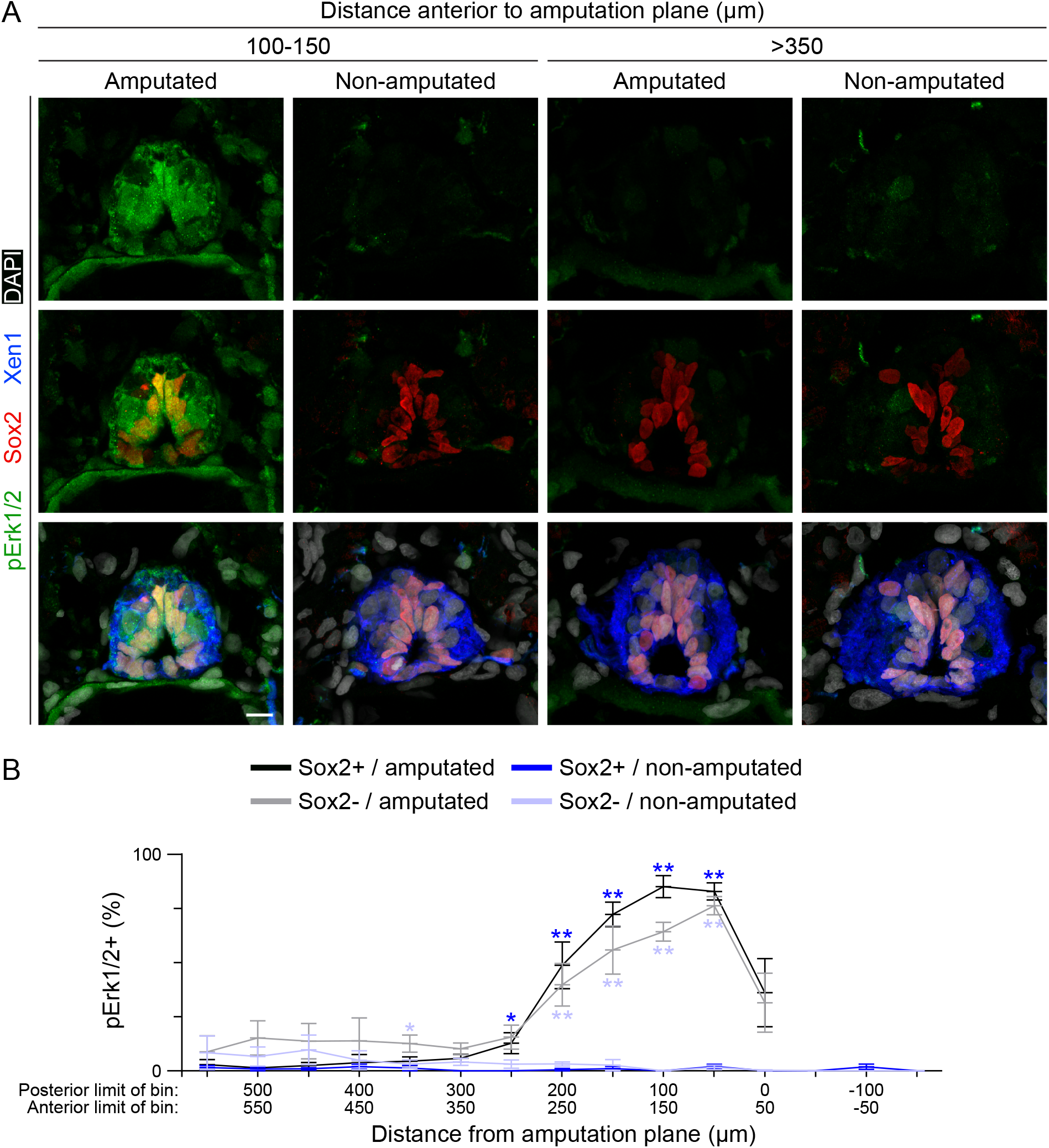
Erk1/2 is activated in spinal cord upon injury. Stage-42 larvae were fixed 20 min post-amputation, then processed for transverse sectioning of the tail and co-immunostaining for Sox2 (neural stem cell), Xen1 (pan neuronal membrane) and pErk1/2 (phosphorylated/activated Erk1/2). Non-amputated siblings were fixed, embedded, and sectioned alongside amputated larvae including up to 105 μm posterior to the plane of amputation. (**A**) Representative maximum-intensity projections of tissue sections at the indicated distance anterior to the plane of amputation. Scale bar is 10 μm. (**B**) In approximately every other tissue section, all cells identified by DAPI and within the spinal cord delineated by Xen1 staining were analyzed for Sox2 and pErk1/2 fluorescent signal above an intensity threshold. Sections were pooled by distance from amputation (0 μm) into 50-μm bins: -x posterior (non-amputated only) and +x anterior. Data are mean±SEM percent of Sox2+ (9-42 cells/section except for amputated samples within 112 μm of the amputation: 0-24 cells/section) and Sox2-(8-44 cells/section) cells that are pErk1/2+. N≥3 independent experiments for total n=4-6 larvae per treatment group. Difference in % pErk1/2+ between amputated and non-amputated larvae was analyzed by ANOVA by bin, and color-coded stars indicate *p<0.05, **p<0.001 in Sox2+ (dark blue) or Sox2-(light blue) cells.

We also found Erk1/2 activation in muscle satellite cells at 1, 2, and 3 days post-amputation (dpa; Figure 2). Examining the spatiotemporal pattern of Erk1/2 activation in Pax7+ cells reveals that from 1 to 3 dpa, signaling is activated primarily in the regenerating tail and up to 100 µm of the stump immediately anterior to the amputation (Figure 2B, C). As regeneration progresses, the proportion of these cells that exhibit active Erk1/2 stays high at the distal end of the regenerate and gradually decreases closer to the stump (Figure 2B, C).

**Figure 2.**
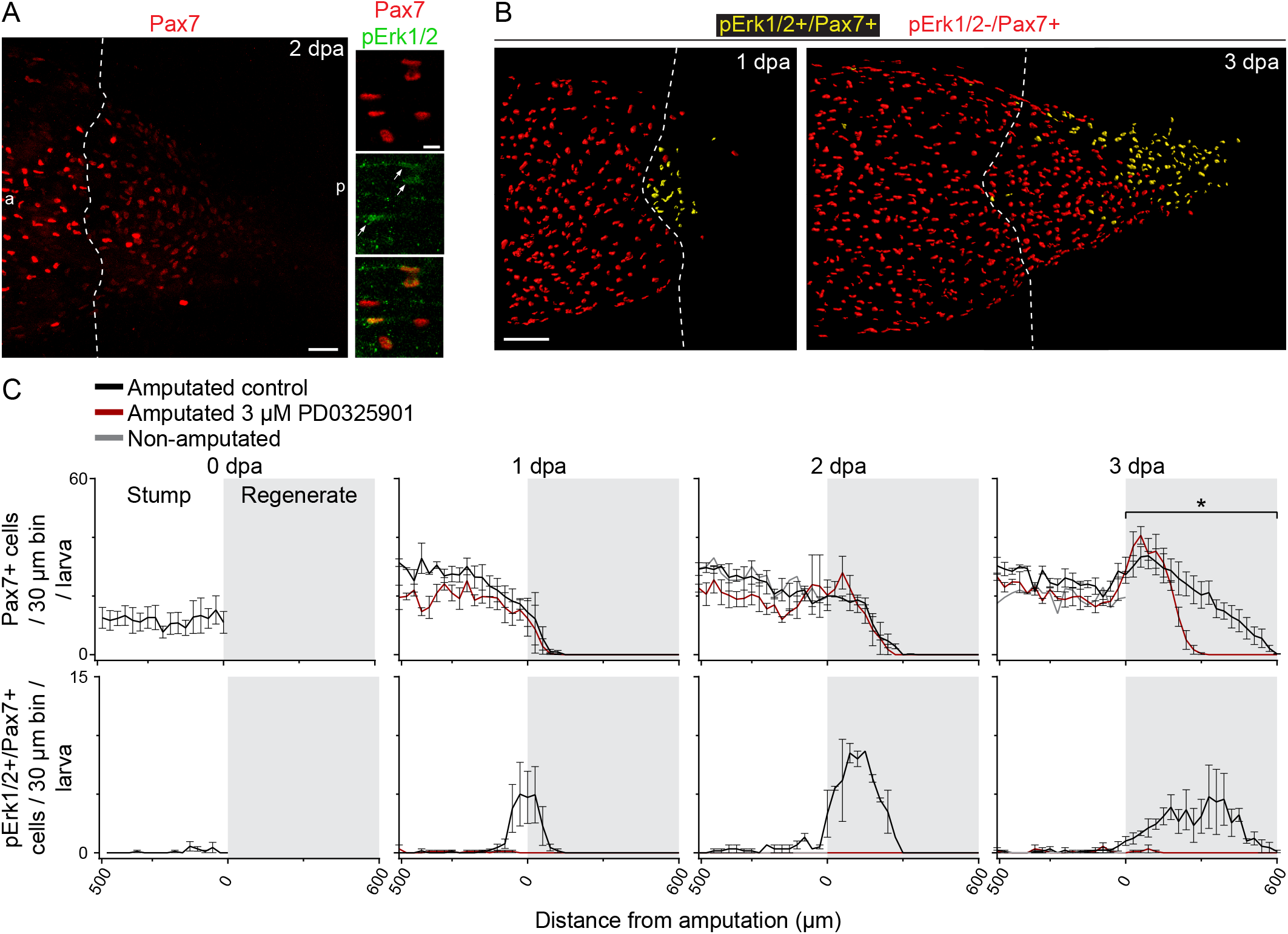
Erk1/2 is activated in muscle satellite cells participating in regeneration. Stage-39 larvae were incubated immediately following amputation with either Mek1/2 inhibitor (3 µM PD0325901) or only vehicle (0.1% DMSO; control) for 1-3 days at 21°C, and then processed for whole-mount Pax7 (muscle satellite cell) and pErk1/2 (activated Erk1/2) immunostaining. (**A**) Representative maximum-intensity projections of larvae at 2 days post-amputation (dpa). Arrows indicate cells that are Pax7+/pErk1/2+. In (**A, B**), dashed lines delineate the border between stump and regenerated muscle. a: anterior, p: posterior. (**B**) Representative volume renderings (Imaris) showing all Pax7+/pErk1/2-nuclei (red) and Pax7+/pErk1/2+ nuclei (yellow). Scale bars are 100 (**A** left, **B**) or 10 (**A** right) µm. (**C**) Data are mean±SEM number of immunopositive cells in each 30 µm-wide region along the longitudinal axis of the tail up to 500 µm anterior to the amputation (0 µm) and the regenerating tail (gray shading). N=3 experiments per time point with n=3 larvae per treatment per experiment. Two-way ANOVA compared the total number of Pax7+ cells in the posterior 500 µm of stump or in the regenerate with or without 3 µM PD0325901; *p<0.05.

Our analysis of Pax7+ cell distribution is consistent with previous findings demonstrating that these cells participate in the regeneration of muscle in the tail (Chen et al., 2006), and additionally shows that there are few Pax7+ cells in the regenerating tissues at 1 dpa which increase in number through 3 dpa (Figure 2C). Pharmacologically inhibiting Erk1/2 signaling by incubating amputated larvae with PD0325901, an inhibitor of the Erk1/2 kinase Mek1/2 (Barrett et al., 2008; Figure 2-supplement 1A), decreases the total number of Pax7+ cells in the regenerated tissue at 3 dpa, while the number of Pax7+ cells in the stump at 1, 2, or 3 dpa, or in the regenerate at 1 or 2 dpa is unaffected (Figure 2C).

Given the activation of Erk1/2 signaling in neural and muscle stem cells upon injury, we investigated whether Erk1/2 signaling is necessary for injury-induced proliferation and found that inhibiting Mek1/2 reduces the number of mitotic cells in all non-fin regenerating tissues (Figure 3A, B), and also reduces the number of mitotic cells in the stump at 1 dpa (Figure 3B). In addition, inhibiting Shp2 phosphatase, which is known to sustain ERK1/2 activation through dephosphorylation of Ras (Bunda et al., 2015), by incubating amputated larvae with SHP099 (Garcia-Fortanet et al., 2016; Figure 2-supplement 1B) for 3 days also dose-dependently reduces the number of mitotic cells in all non-fin regenerating tissues and specifically in the spinal cord (Sox2+), but does not affect the number of mitotic cells in the stump (Figure 3A, C).

**Figure 3.**
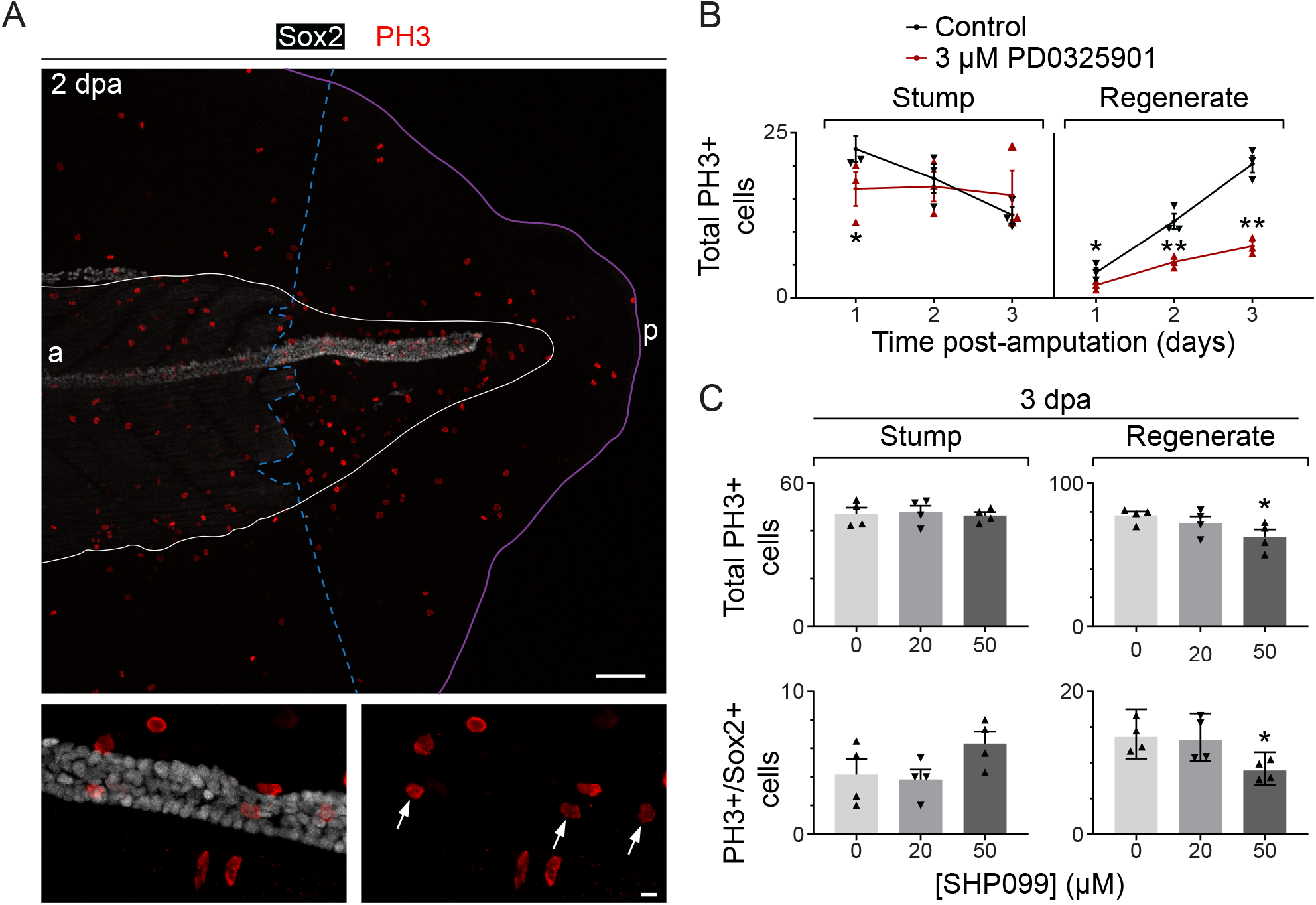
Erk1/2 signaling is important for cell proliferation in the regenerating tail. Stage-39 larvae were incubated immediately following amputation with 3 µM PD0325901, 20 or 50 µM SHP099, or only vehicle (0.1 or 0.5% DMSO; control) for 1-3 days at 21°C, and then processed for whole-mount immunostaining for PH3 (mitotic cell) and Sox2 (neural stem cell). (**A**) Representative maximum-intensity projections. Dashed, blue line delineates the border between stump and regenerated muscle. White line delineates analyzed region. Purple line shows the outside edge of the fin; a: anterior, p: posterior. Scale bars are 100 (top) or 10 (bottom) µm. Arrows indicate cells that are immunopositive for PH3 and Sox2. (**B, C**) Data are mean±SEM (**B**), mean+SEM (**C**) or geometric mean±95% CI (**C** lower right) number of PH3+ cells (**B, C**) or PH3+/Sox2+ cells (**C**) in the regenerating tail (regenerate) and in the first 300-500 µm anterior to the amputation (stump), excluding the fin. (**B, C**) Triangles show means (**B, C**) or geometric means (**C** lower right) from independent experiments. For each timepoint, N=3-4 experiments with n=3-6 larvae per treatment per experiment. Statistical analyses were performed by two-way ANOVA (**B, C**) followed by Dunnett’s means comparison (**C**); *p<0.05, **p<0.001.

Altogether, these data suggest that, upon injury, Erk1/2 is activated in neural and muscle stem cells to promote their proliferation in regenerating tissues.

### Erk1/2 signaling is important for regeneration of spinal cord and muscle

We then measured the extent of spinal cord, skeletal muscle, and notochord regeneration (Figure 4-supplement 1) in response to pharmacological inhibition of Erk1/2 signaling and found a dose-dependent reduction (Figure 4 and Figure 4-supplement 2). Results show that inhibition of Erk1/2 activation with the Mek1/2 inhibitor PD0325901 (Figure 4A-C), or the Shp2 phosphatase inhibitor SHP099 (Figure 4A, D, E), reduces both spinal cord and muscle regeneration in addition to notochord (Figure 4-supplement 2). Similarly, inhibition of Mek1/2 with the inhibitor PD98059 also reduces muscle regeneration (Figure 4-supplement 3).

**Figure 4.**
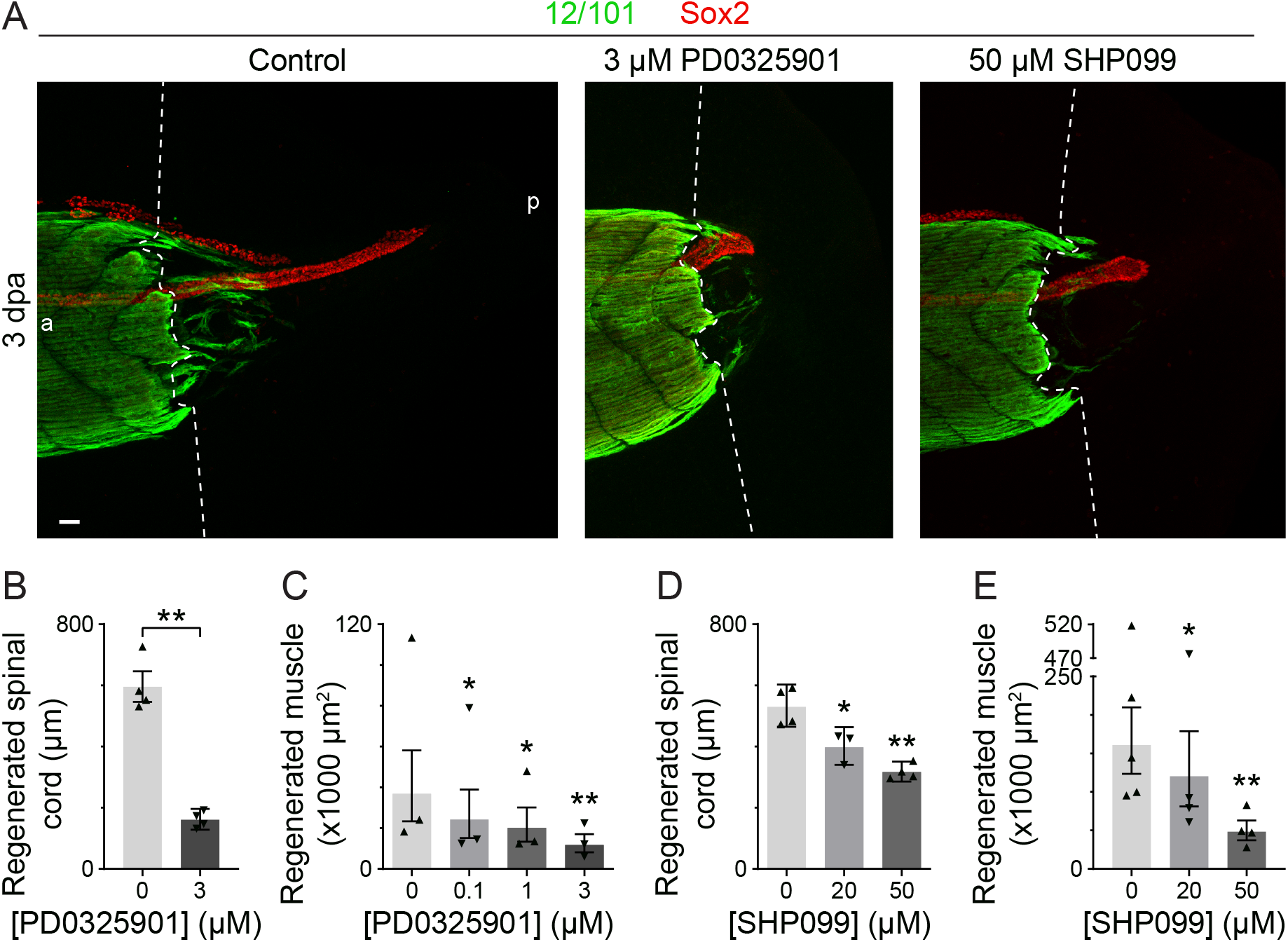
Activation of the Erk1/2 signaling pathway is important for muscle and spinal cord regeneration. Stage-39 larvae were incubated immediately following amputation with Mek1/2 inhibitor (0.1-3 µM PD0325901), Shp2 inhibitor (20 or 50 µM SHP099) or only vehicle (0.1 or 0.5% DMSO; control; 0 µM) and then grown for 3 days at 21°C. Larvae were processed for 12/101 (skeletal muscle) and Sox2 (neural stem cell) whole-mount immunostaining, and then imaged with a confocal microscope. (**A**) Representative maximum-intensity projections (MIPs). Dashed lines delineate the border between stump and regenerated muscle; a: anterior, p: posterior; scale bar is 50 µm. (**B**-**E**) Data are back-transformed mean±95% CI length of the regenerated spinal cord stained by Sox2 and measured in MIPs of the regenerating tail (**B, D**) or geometric mean±95% CI sum of the area stained by 12/101 in the regenerating tail measured in each frame of the z-stack (**C, E**). Triangles show back-transformed (**B, D**) or geometric (**C, D**) means from independent experiments. (**B**-**E**) N=3-4 experiments with n=3-6 larvae per treatment per experiment. Statistical analyses were performed by two-way ANOVA (**B**-**E**) followed by Dunnett’s means comparison (**C**-**E**); *p<0.05, **p<0.001.

Genetically inhibiting Erk1/2 signaling by knocking down expression of *map2k1.L*, the gene encoding the single MEK1/2 homolog in *Xenopus laevis,* during regeneration using a UV-activated, translation-blocking morpholino targeting *map2k1.L* mRNA (*map2k1.L*-mo; Figure 5-supplement 1A-D) mimics the effect of pharmacologically inhibiting Mek1/2 signaling by reducing regeneration of spinal cord, muscle (Figure 5A-C) and notochord (Figure 5-supplement 2A, B).

**Figure 5.**
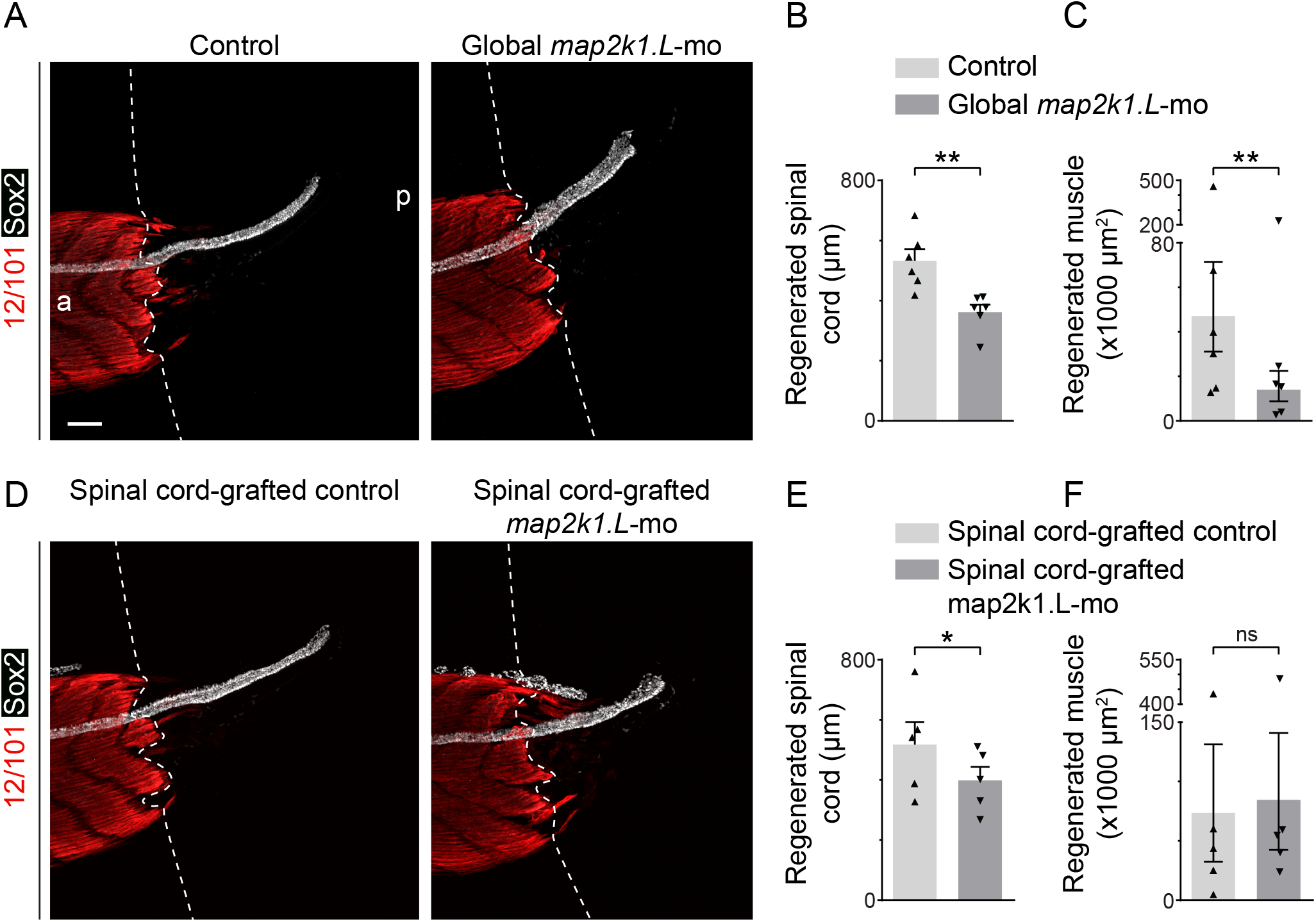
Tissue-specific requirement for Erk1/2 activation for spinal cord and muscle regeneration. One-cell-stage embryos were injected with 9-11 pmol *map2k1.L*-targeted morpholino (*map2k1.L*-mo) mixed with 11-13 pmol complementary, caging photo-morpholino. Animals were grown in the dark, exposed to UV for 3 min 4-6 h prior to amputation at stage 39 to uncage the *map2k1.L*-mo, and then allowed to regenerate for 3 days in the dark at 21°C. (**A, D**) Images are representative maximum-intensity projections (MIPs) with dashed lines delineating the border between stump and regenerated muscle; a: anterior, p: posterior; scale bar is 100 µm. (**A**-**C**) Controls are WT siblings exposed to UV and grown under the same conditions. (**D**-**F**) Larvae with *map2k1.L*-mo only in the spinal cord were generated by replacing the neural plate of stage 12.5-13 WT embryos with neural plate from their siblings with global *map2k1.L*-mo. Controls were generated by grafting neural plate from WT siblings onto WT embryos which were then exposed to UV and grown under the same conditions. (**B, C, E, F**) Data are mean+SEM length of regenerated spinal cord stained by Sox2 (neural stem cell) and measured in MIPs of the regenerating tail (**B, E**) or back-transformed mean±95% CI sum of the area stained by 12/101 (mature skeletal muscle) in the regenerating tail and measured in each frame of the z-stack (**C, F**). Scattered triangles represent means (**B, E**) or back-transformed means (**C, F**) of independent experiments. N=5-6 experiments with n=3-9 larvae per treatment per experiment. Statistical analyses were performed by two-way ANOVA; *p<0.05, **p<0.001.

To assess the tissue-specific necessity of Erk1/2 signaling for regeneration, we generated larvae with *map2k1.L* knockdown only in spinal cord by replacing the neural plate of stage 12.5-13 wild-type embryos with neural plate from siblings with *map2k1.L*-mo in all tissues, and growing them to larval stages (Figure 5-supplement 1E). This spinal cord-specific knockdown decreases spinal cord (Figure 5D, E) and notochord (Figure 5-supplement 2C, D) regeneration, but does not significantly affect muscle regeneration (Figure 5D, F), suggesting that the dependence of muscle regeneration on Erk1/2 activity is independent of this signaling in the spinal cord.

### Injury induces an increase in Ca^2+^ dynamics that persists in regenerating tissues

Previous studies in our lab showed that during development, inhibition of Erk1/2 signaling decreases spontaneous Ca^2+^ activity in embryonic spinal cord neurons (Swapna & Borodinsky, 2012) and that interfering with Ca^2+^ release from internal stores alters the proliferation of myogenic progenitors in the regenerating larval tail (Tu & Borodinsky, 2014), suggesting that Ca^2+^ signaling may interact with Erk1/2 signaling during regeneration. To investigate Ca^2+^ activity in regenerating tissues *in vivo*, the tails of stage-39 larvae expressing the genetically-encoded Ca^2+^ sensor GCaMP6s were time-lapse-imaged before and after amputation. The tail was imaged laterally and, because of the opacity of the tissue, the cells visualized using this method were primarily those lateral to the axial musculature where muscle progenitors reside. Amputation increases the total number of Ca^2+^ transients and active cells throughout an 800 µm-long region of tail just anterior to the amputation relative to sham siblings which were subjected to the same anesthetic and embedding protocols as other treatments but were not amputated (Figure 6A-C and Figure 6-Videos 1 and 2). An increase in frequency of Ca^2+^ transients per active cell (mean±SEM increase of 1.2±0.29 transients/15 min/cell, n=8 larvae, p = 0.0475 relative to sham) contributes to this overall increase in Ca^2+^ activity. Analysis of the spatial distribution of Ca^2+^ activity in relation to the site of injury shows no differences between any 100 μm-wide regions up to 800 µm from the amputation (p = 0.1206, n = 9 larvae), indicating that distant and adjacent cells show a similar increase in Ca^2+^ dynamics following injury.

**Figure 6.**
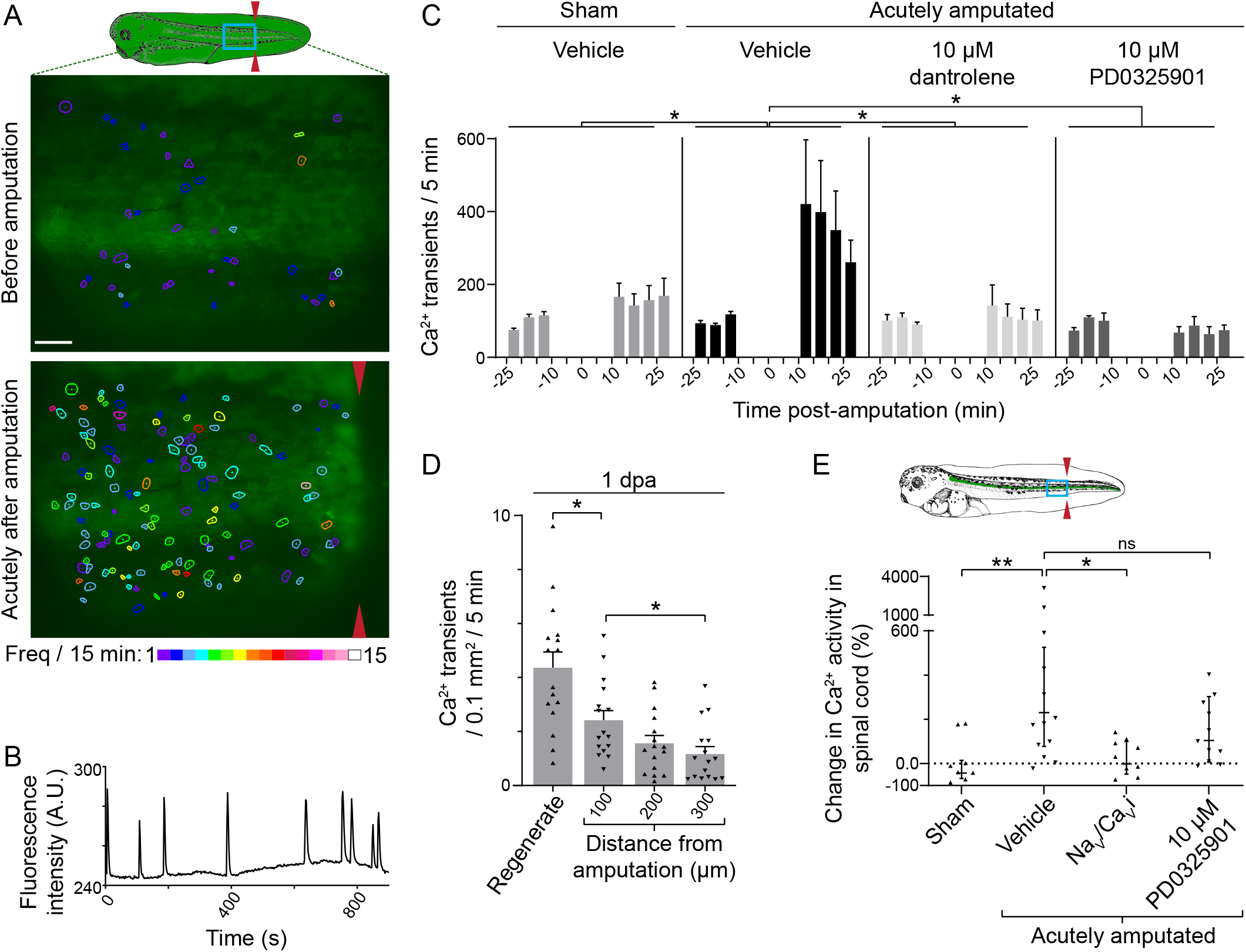
Injury enhances Ca^2+^ activity that persists in regenerating tissues. (**A**-**C**) Stage-39 larvae globally expressing GCaMP6s were time-lapse-imaged before and after amputation using a fluorescent stereoscope at 0.5 Hz. The number of transients lateral to the myotome and up to 800 µm from the amputation site were counted during 25 to 10 min before and 10 to 30 min following the amputation. Immobilized larvae were incubated with indicated drugs or only vehicle starting just after the first imaging period, 10 min prior to amputation (t=0). Sham experienced the same embedding and anesthesia procedures as the other groups but were not amputated. (**A** top) Schematic of a stage-39 larva (illustration adapted from Nieuwkoop and Faber, 1994) showing the region imaged (blue box). (**A** middle, bottom) Images are from a representative larva before and after amputation (red arrowheads). Cells exhibiting Ca^2+^ transients are outlined with colors corresponding to transient frequency per 15 min. (**B**) Example trace of a single cell showing multiple Ca^2+^ transients. (**C**) Data are mean+SEM number of Ca^2+^ transients per 5 min, n=6-9 larvae per treatment. Statistical analysis was performed by ANOVA comparing the change in total number of transients during the first 15 min of imaging before and after amputation in each larva, followed by Dunnett’s means comparison to vehicle-only amputated larvae. (**D**) Larvae (n=12) globally expressing GCaMP6s were amputated at stage 39, grown at 21°C for 22-26 hours, and then embedded and imaged as described above for 10 min. Data are mean+SEM number of Ca^2+^ transients per 0.1 mm^2^ of tail per 5 min. Scattered triangles in (**D, E**) represent values for individual larvae. (**E**) Larvae expressing GCaMP6s only in the spinal cord were generated by replacing the neural plate of stage 13-14 WT embryos with neural plate from their siblings globally expressing GCaMP6s. The resulting larvae were time-lapse imaged at stage 42 from 30 to 10 min before and 10 to 50 min following amputation at 0.5 Hz. The schematic shows a stage-42 larva with the region imaged delineated by the blue box. Red arrows mark the site of amputation. Larvae were incubated with indicated drugs or only vehicle starting 10 min prior to amputation. Na_V_/Ca_V_i is a mixture of voltage-gated Na^+^ and Ca^2+^ channel blockers: 1 µM GIVA ω-conotoxin, 20 nM calcicludine, 1 µM flunarizine, and 1 µg/ml tetrodotoxin. Sham larvae experienced the same embedding and anesthesia procedures as other groups but were not amputated. For each larva, Ca^2+^ transients in the region of spinal cord that was visible during both imaging periods were counted. Data are geometric mean±95%CI percent change in the number of Ca^2+^ transients per 5 min before and after the amputation for each larva, n=8-13 larvae per treatment. In (**D, E**) statistical analyses were performed by ANOVA followed by Tukey’s means comparison (**D**) or Dunnett’s means comparison to vehicle-only amputated larvae (**E**). In (**C**-**E**), *p<0.05, **p<0.001.

Previous studies have implicated Ca^2+^ release from intracellular stores during embryonic myogenesis (Ferrari & Spitzer, 1999). Consistent with development, we find that the ryanodine receptor blocker dantrolene significantly suppresses the amputation-induced increase in Ca^2+^ activity (Figure 6C), demonstrating that Ca^2+^ release contributes to injury-induced Ca^2+^ transients.

To examine the relationship between Ca^2+^ and Erk1/2 signaling, we inhibited Mek1/2 with PD0325901 and found that this prevents the injury-induced increase in Ca^2+^ activity in cells lateral to the axial musculature (Figure 6C), suggesting that in these cells, injury-induced Erk1/2 activation recruits Ca^2+^ activity to promote regeneration of the larval tail. To determine if, like Erk1/2 signaling, Ca^2+^ activity continues as regeneration progresses, we assessed Ca^2+^ dynamics in the regenerating tail at 1 dpa and found that Ca^2+^ transients persist in regenerating tissues and in the adjacent, posterior region of the stump (Figure 6D and Figure 6 – Video 3). In contrast to acutely-amputated larvae, at 1 dpa Ca^2+^ activity in the stump inversely correlates with the distance from the amputation, with regenerated tissues exhibiting significantly higher Ca^2+^ activity than the stump (Figure 6D).

To assess Ca^2+^ activity specifically in the regenerating spinal cord, larvae expressing GCaMP6s only in spinal cord were generated by replacing the neural plate of stage 13-14 wild-type embryos with neural plate from siblings expressing GCaMP6s and growing them to larval stage 42. Approximately 300 µm of spinal cord anterior to the amputation was visualized through the surrounding tissues. Data show that amputation induces an acute increase in Ca^2+^ transient frequency in the spinal cord that persists for at least 50 min following injury (Figure 6E and Figure 6 – Videos 4 and 5). During embryonic spinal cord development, neuronal Ca^2+^ transients are dependent on both Ca^2+^ influx and release from internal stores (Gu et al., 1994; Gu & Spitzer, 1995). We find that inhibiting voltage-gated Na^+^ and Ca^2+^ channel blockers (Borodinsky et al., 2004) suppresses the injury-induced increase in Ca^2+^ transients in the spinal cord (Figure 6E), indicating that these transients are dependent on Ca^2+^ influx.

In contrast to tissues associated with muscle (Figure 6C), Mek1/2 inhibition did not significantly impact the injury-induced increase in Ca^2+^ activity in the spinal cord (Figure 6E), suggesting that the relationship between Erk1/2 and Ca^2+^ signaling may be distinct in these two tissues during regeneration.

## DISCUSSION

Promoting regeneration in species with limited inherent regenerative capacity may depend on identification of differences in response to injury between regeneration-proficient and regeneration-limited organisms. These include changes in both the stem cell niche and in intracellular signaling in stem cells that generate new tissue. Previous studies show that recruitment of ERK1/2 signaling in diverse *in vitro* and *in vivo* models is an important regulator of muscle satellite cell function. In mouse muscle satellite cells, FGF-induced ERK1/2 activation is necessary to promote cell cycle progression both *in vitro* (Jones et al., 2001) and *in vivo* (Griger et al., 2017), and hyper-activation of ERK1/2 signaling in muscle satellite cells prevents them from returning to quiescence (Shea et al., 2010). However, the precise spatiotemporal pattern of ERK1/2 activation to promote regeneration, and its role in other tissues have not been previously characterized. Our study demonstrates that Erk1/2 activation in muscle and neural stem cells in response to injury is necessary for cell proliferation and regeneration of muscle and spinal cord in *Xenopus laevis* larvae. We find that in addition to initial activation by Mek1/2, sustained Erk1/2 signaling supported by Shp2 action is also important for spinal cord and skeletal muscle regeneration. This is consistent with the previously reported necessity of SHP2 signaling for activation of mouse adult muscle satellite cells but not fetal myoblasts (Griger et al., 2017), suggesting that the signaling mechanisms are conserved between adult mice and *Xenopus* larvae.

During amphibian spinal cord regeneration, regenerated neural tissue is primarily derived from neural stem cells adjacent to the injury (Lin et al., 2007; Mchedlishvili et al., 2007; Sabin et al., 2015). Our findings suggest that in the injured spinal cord, Erk1/2 is activated specifically in these cells that are most likely to contribute to its regeneration. Similarly, this study demonstrates that in muscle satellite cells, Erk1/2 signaling is specifically activated in the regenerate and in the stump adjacent to the amputation. As regeneration progresses, Erk1/2 activation in muscle satellite cells decreases adjacent to the amputation, where muscle is often differentiating, but remains high in the distal tip which is usually in advance of muscle differentiation. Altogether, these findings suggest that Erk1/2 activation is selecting a subpopulation of stem cells to proliferate and contribute to the regenerating muscle and spinal cord. In addition, spinal cord-specific reduction in Erk1/2 signaling mimicked the effects of global inhibition on spinal cord and notochord regeneration, but not muscle regeneration, suggesting that Erk1/2 activation may be independently necessary in spinal cord and muscle for proficient regeneration. This study suggests that activation of Erk1/2 signaling in both neural and muscle stem cells is necessary to trigger their respective regenerative potential.

We show *in vivo* that an acute, amputation-induced increase in Ca^2+^ signaling in both spinal cord and tissues lateral to the axial musculature is independent of distance from the injury, while at 1 day post-amputation, the frequency of Ca^2+^ transients inversely correlates with the distance from the regenerating tissues and directly with the spatial pattern of active Erk1/2 signaling in muscle satellite cells. This difference could indicate there are two distinct phases of Ca^2+^ signaling initiated by the injury: an immediate but temporary increase that spans most of the remaining tail, and a persistent increase specific to the cells contributing to the regenerate. In addition, we find that Mek1/2 blockade prevents the acute, amputation-induced Ca^2+^ dynamics in non-neural tissues but not in the spinal cord, despite the fact that Erk1/2 is induced in spinal cord during this time, demonstrating that diversity in pathway crosstalk may distinguish Ca^2+^ signaling in these two tissues. Previous studies show that Ca^2+^ signaling results in divergent outcomes depending on the precise spatiotemporal dynamics (Samanta & Parekh, 2017; Wacquier et al., 2019), therefore the distinct patterns of injury-induced Ca^2+^ activity that we identified could induce different responses.

In regenerating *Xenopus laevis* larval tails, the spatial patterns of both Erk1/2 activation and increased Ca^2+^ activity that we observed correlate with the previously reported patterns of expression of components of Fgf (Lin & Slack, 2008; Love et al., 2013) and Wnt signaling pathways (Lin & Slack, 2008), in addition to active ROS signaling (Love et al., 2013). Moreover, Bmp (Lin & Slack, 2008), Wnt (Lin et al., 2012; Lin & Slack, 2008), and ROS (Love et al., 2013) signaling are all upstream of the Fgf signaling that is necessary for *Xenopus* larval tail regeneration (Lin et al., 2012; Lin & Slack, 2008; Love et al., 2013). During embryonic development in *Xenopus laevis*, Erk1/2 activation is dependent on Fgf signaling (Christen & Slack, 1999), and since regeneration often reiterates developmental pathways (Cardozo et al., 2017), Erk1/2 signaling may be activated by these ROS and Bmp/Wnt/Fgf signaling networks, which then activate Ca^2+^ signaling in muscle progenitors. Non-canonical hedgehog signaling is another candidate for initiating Ca^2+^ activity in response to injury because during development it promotes Ca^2+^ activity in the embryonic spinal cord (Belgacem & Borodinsky, 2011), and is necessary for spinal cord and skeletal muscle regeneration following tail amputation in *Xenopus laevis* larvae (Hamilton et al., 2021). In addition, Hippo signaling mediated by Yap1 is also important for *Xenopus laevis* larval tail regeneration (Hayashi et al., 2014), and is important for EGFR-mediated Erk1/2 activation in regenerating *Xenopus* retina (Hamon et al., 2019), suggesting it as another candidate for mediating Erk1/2 during regeneration. Future investigation elucidating the interaction between these pathways and Erk1/2-Ca^2+^ signaling has the potential to unify our understanding of the signaling that is driving regeneration into a single network, and could be the next step towards gaining therapeutic control of muscle and neural stem cell status to promote regeneration in humans.

## MATERIALS AND METHODS

### Key resources table

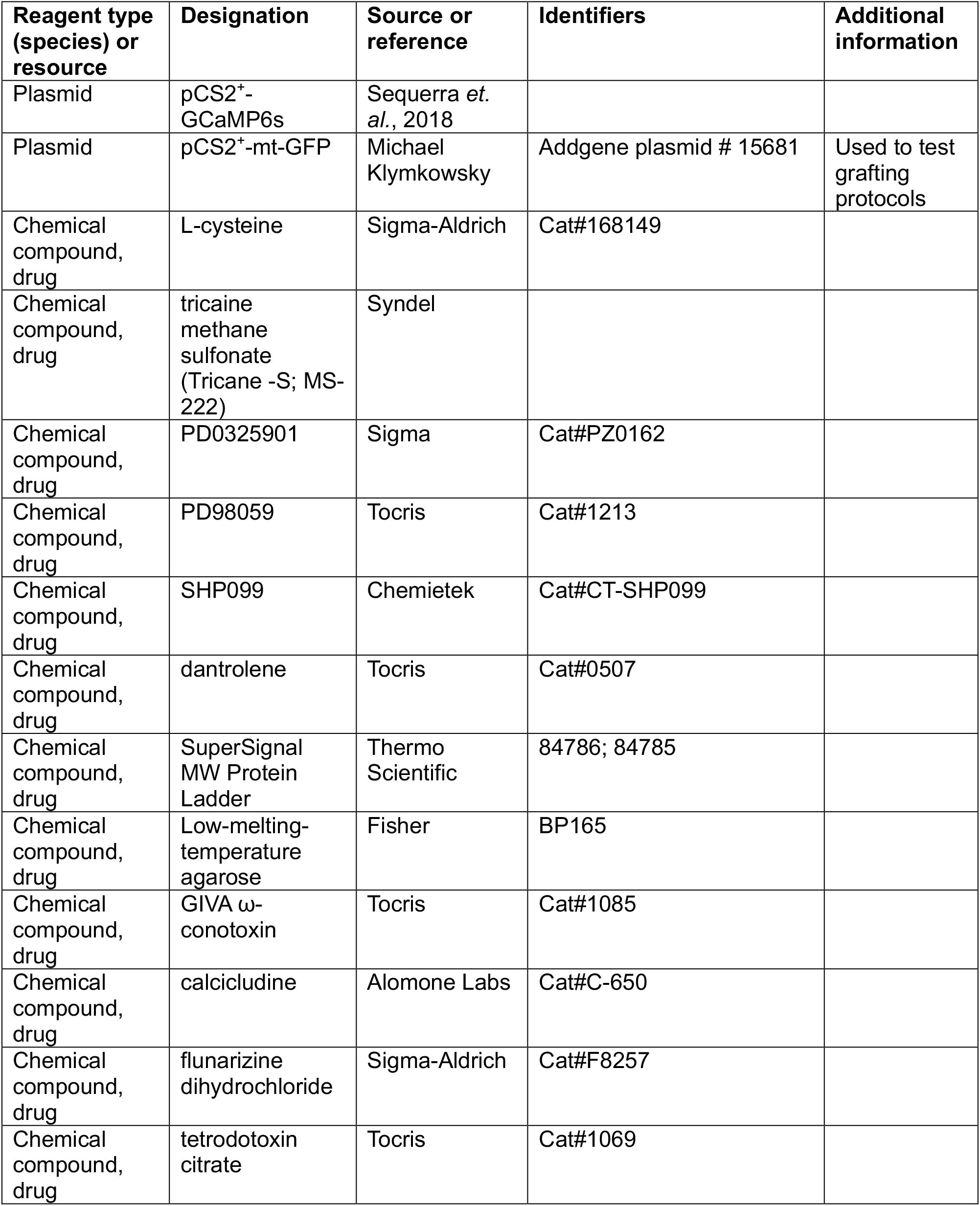
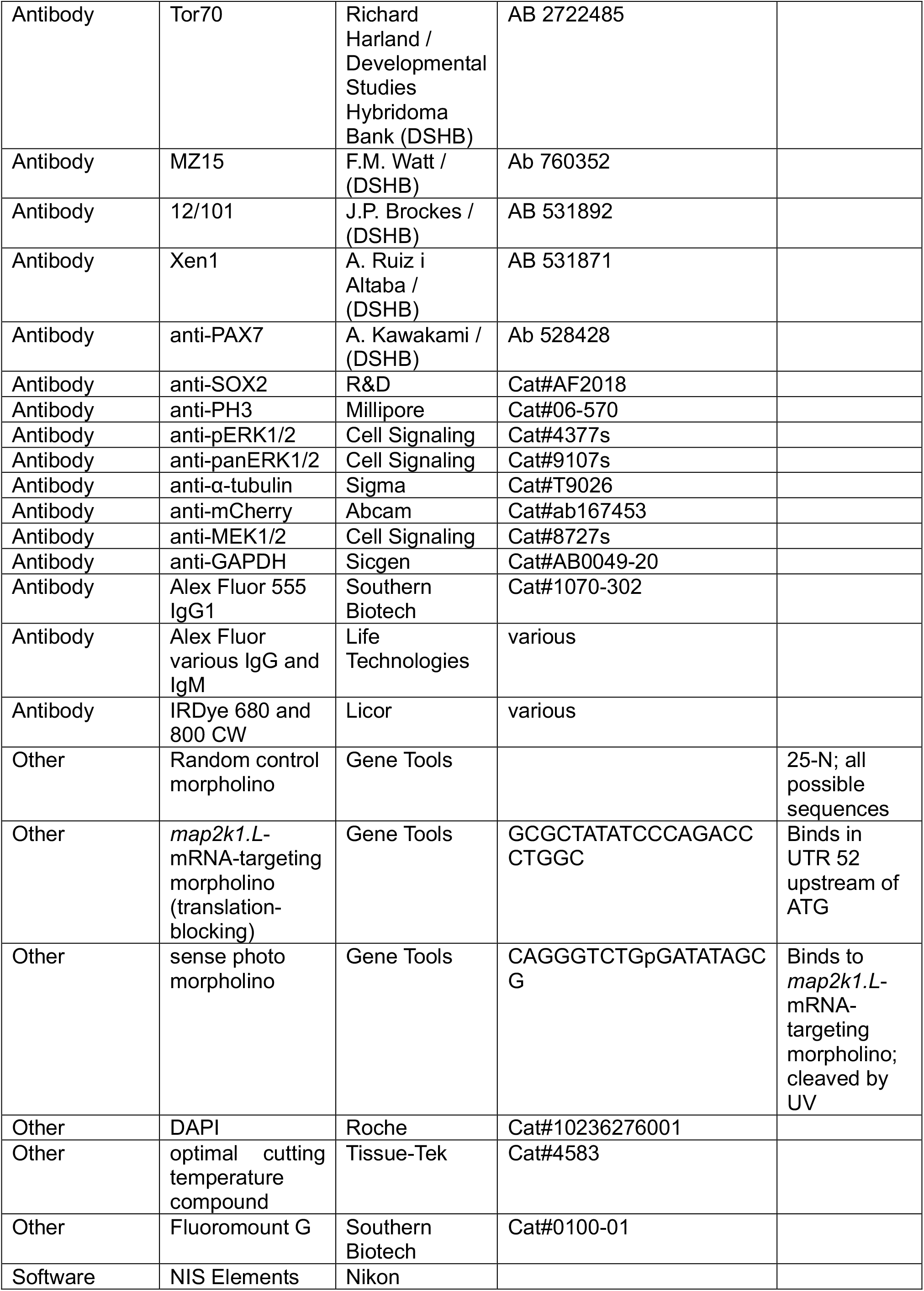
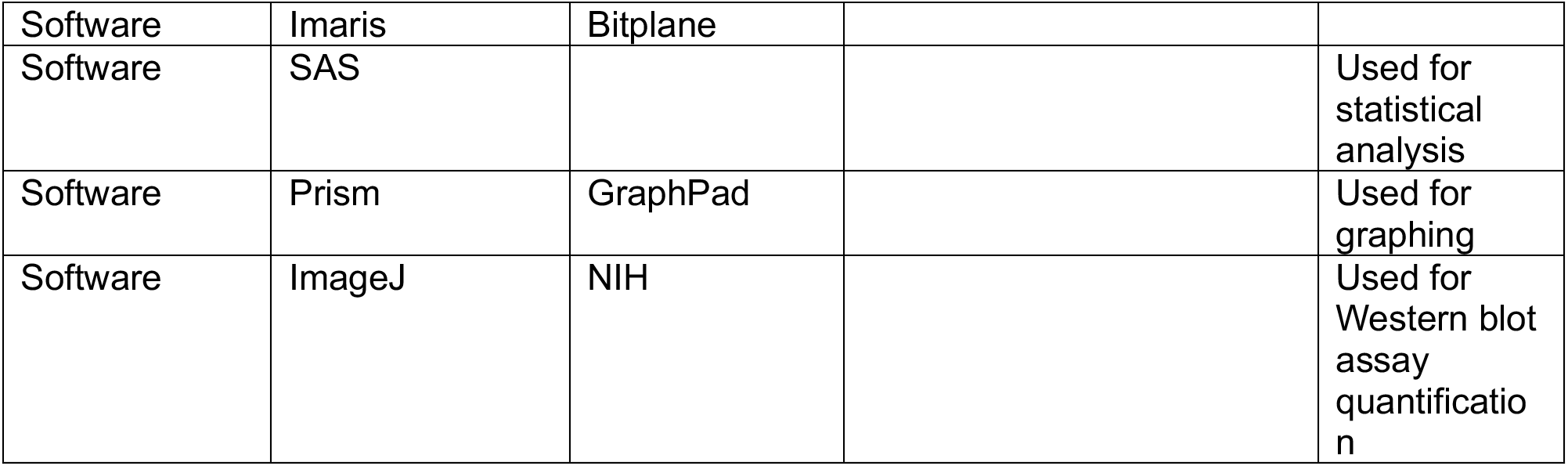

#### Animals

Embryos were generated by *in vitro* fertilization as previously described (Sive et al., 2000). Fertilized embryos were either partially dejellied in 2% cysteine (pH 8) for microinjection of constructs at 1-4-cell stages or left in the full jelly coat. Animals were handled humanely according to IACUC guidelines and an approved animal protocol. Embryos and larvae were grown in 0.1X Marc’s Modified Ringer’s (MMR) solution in mM: 10 NaCl, 0.2 KCl, 0.1 MgSO_4_, 0.5 HEPES, 0.01 EDTA, and 0.2 CaCl_2_.

#### Tail amputation and drug application

Amputations were performed at stage 39 or 42 (Nieuwkoop & Faber, 1994; 56 or 80 h post-fertilization) with a surgical scalpel at the midpoint of the tail (approximately 23 chevrons from the head) after anesthetizing larvae for 10 min in 0.02% tricaine methanesulfonate (TMS) in 0.1X MMR with 5 mM HEPES, pH 7.4. Larvae were then washed with 0.1X MMR several times before transferring to final solution which included either the indicated drug, or vehicle only. While regenerating, larvae were grown in 24-well plates at a density of 3 larvae in 2 ml of solution per well. Solutions were replaced daily. All drugs were premixed into MMR and added immediately following amputation with the exception of the live-imaging experiments where drugs were added with the anesthetic 10 min prior to amputation to allow sufficient time for drug access to agarose-mounted larva before imaging.

#### Immunostaining in tissue sections

Larvae were fixed overnight at 4°C in fresh, 4% paraformaldehyde. They were rinsed briefly with phosphate-buffered saline (PBS), then dehydrated in 15-30% sucrose until they sank and embedded in optimal cutting temperature compound. Amputated larvae and their non-amputated siblings were arranged side-by-side in each block, partially frozen in liquid nitrogen, then completely frozen and stored at -20°C, and transversely sectioned 14-15 µm thick. Slides were probed with antibodies against SOX2 1:100 (R&D AF2018), pERK1/2 1:50 (Cell Signaling 4377s), and Xen1 1:100 (deposited to Developmental Studies Hybridoma Bank (DSHB) by Ruiz i Altaba, A.) overnight at 4°C. Slides were incubated with secondary antibodies for 2 h at room temperature followed by DAPI, then rinsed and mounted. Sections were imaged on a Nikon A1 or C2 confocal microscope with a 100X objective and z-steps of 1 µm. Images were analyzed using NIS Elements where nuclei in the spinal cord, demarcated by Xen1 staining, were counted as Sox2 and/or pErk1/2 immunopositive if at least half of the nucleus was above an intensity threshold between 1.5 and 3x background for each immunostaining.

#### Whole-mount immunostaining

During regeneration, larvae were anesthetized for 5 min in 0.02% TMS, then fixed overnight at 4°C in MEMFA salt solution (100 mM MOPS pH 7.4, 2 mM EGTA pH 8.0, 1 mM MgSO_4_, sterilized with a 0.22-µm filter) and 3.7% formaldehyde. Samples were rinsed briefly in PBS, bleached overnight in 1:2 H_2_O_2_:Dent’s fixative, and then permeabilized for at least 5 h with 5 washes of 0.5% Triton-X100 in PBS (PBT). Samples were blocked in 10% goat serum (GS) or 2-10% bovine serum albumin (BSA) in 0.5% PBT, and then incubated with primary antibody overnight at 4°C in 0.1% PBT with 10% GS or 2-10% BSA. Antibodies used included 12/101s 1:50 (deposited to DSHB by Brockes, J.P.), MZ15 1:100 (deposited to DSHB by Watt, F.M.), PAX7 1:50 (deposited to DSHB by Kawakami, A.), Tor70 1:4 (deposited to DSHB by Harland, R.), Xen1 (DSHB) 1:100, pERK1/2 1:50 (Cell Signaling 4377s), SOX2 1:100 (R&D AF2018), phospho-histone-H3 1:300 (PH3; Millipore 06-570). Samples were then washed in 0.5% PBT for 5-7 h and then incubated with secondary antibody (1:300) in 0.1% PBT with 10% GS or 2-10% BSA overnight at 4°C. They were washed in 0.5% PBT for 5-7 hours, then heads were removed and tails were mounted in 80% glycerol. They were imaged on a Nikon A1 or C2 confocal microscope with a 10 or 20X objective. 12/101, an antibody targeted against a sarcoplasmic reticulum membrane protein of mature muscle cells, was imaged with 5 or 7 µm z-steps while nuclear stainings (Sox2, Pax7) were imaged with 3 or 5 µm z-steps. Images were analyzed using NIS Elements (12/101-immunolabeled area above intensity threshold per z-step; MZ15-, Tor70- or Sox2-immunolabeled length; Nikon Inc.) or Imaris (PH3, Pax7, pErk1/2-immunopositive nuclei count; Bitplane Inc.) software.

To measure the extent of regeneration, the stump was distinguished from the regenerating region using a combination of specific staining and tissue morphology. Sox2 (neural stem cell) and Xen1 (pan-neuronal membrane) immunostainings show the same extension of regenerated spinal cord through at least 3 dpa, supporting the use of Sox2 immunolabeling to report the extent of spinal cord regeneration through 3 dpa (Figure 4-supplement 1A). Regenerated muscle was distinguished from original muscle by the lack of chevron organization that persists through 5 dpa (Figure 4-supplement 1B, C), and this is not affected by treatment with the Mek1/2 inhibitor PD98059 (Figure 4-supplement 1B). We found that Tor70, which has been previously reported to stain immature notochord in *Xenopus* embryos only up to stage 28 (Bolce et al., 1992), also specifically stains regenerating notochord through 4 days post amputation (dpa) at stage 39 (Figure 4-supplement 1D). We predominantly assessed regeneration at 3 dpa because at this time point wild-type (WT) larvae exhibit significant spinal cord and muscle regeneration (Figure 4-supplement 1E).

#### Mek1/2 knockdown

*Xenopus laevis* only has a single MEK1/2 homolog gene designated *map2k1*. The efficacy of *Map2k1.L*-targeted morpholino (GCGCTATATCCCAGACCCTGGC; *map2k1.L*-mo) knockdown was assessed relative to random control morpholino (25-N; all possible sequences) by Western blot assay of animals at stage 22, 29, 32 or 39 that had been injected at 1-cell-stage with 9 pmol of morpholino (Figure 5-supplement1A, B).

For all other experiments, to prevent premature knockdown during early development, *map2k1.L*-mo was premixed with complementary, photo morpholino (CAGGGTCTGpGATATAGCG; photo-mo) at a ratio of 1:1.2 while protected from UV light (white light with a yellow filter; Dolan-Jenner 68600903566). This mixture was heated to 65°C for 20 min, then cooled and 20-24 pmol total morpholino was injected into 1-cell-stage embryos. Animals were grown in the dark except for a 3-min exposure to UV light using an LED array-based UV (365) light source (Gene Tools, LLC.) 4-6 h prior to amputation at stage 39 to cleave the photo-mo and promote its disassociation from the *map2k1.L*-mo, leading to *map2k1.L*-mo activation. Larvae were then allowed to regenerate for 3 days in the dark at 21°C before fixation for whole-mount immunostaining.

#### Western blot assays

Three larvae per sample were snap-frozen in liquid nitrogen, then homogenized at room temperature with a 1 ml glass pestle in 50 μl/larva room-temperature RIPA buffer containing 50mM Tris (pH 7.5), 150 mM NaCl, 0.5% NP-40, 0.05% SDS, 0.002% EDTA, protease inhibitor cocktail (Thermo Fisher Scientific), and phosphatase inhibitor cocktail (Sigma), then flash-frozen again in an Eppendorf tube and stored at -80°C. Samples were thawed on ice, spun at 16,100 RPM for 10-12 min, and the pellet and fatty layer were discarded. The remaining supernatant was mixed with Laemmli buffer and boiled for 5 min. Samples and SuperSignal MW Protein Ladder were then run on a 10% SDS-PAGE gel followed by overnight transfer to PVDF membrane at 4°C. Transfer was confirmed with Ponceau staining and blots were immunostained for fluorescent detection on Odyssey CLx (Licor). Primary antibodies used were alpha-tubulin 1:500 (Sigma T9026), MEK1/2 1:1000 (Cell Signaling 8727s), and GAPDH 1:10,000 (Sicgen AB0049-20) and were incubated for 1 h at room temperature. Secondary antibodies were used at 1:10,000 (Licor) and incubated for 10 min using SNAPid 2.0 (Millipore). ImageJ was used to calculate relative protein levels in immunoblots.

#### Neural tissue grafting

To obtain spinal cord-specific GCaMP6s expression or *map2k1.L* knockdown, we replaced the posterior neural plate of wild-type (WT) embryos at stages 12.5-14 with neural plate dissected from sibling embryos expressing GCaMP6s or containing *map2k1.L*-morpholino, respectively. The grafting technique was similar to that previously described (Gargioli & Slack, 2004), with the following modifications: grafting was performed on a dish with a plastic grid on the bottom to restrain embryos from rolling; no enzymes or drugs were used to promote tissue separation; a Ringer’s solution (4.6 mM Tris base; 116 mM NaCl; 0.67 mM KCl; 1.3 mM MgSO_4_; 2 mM CaCl_2_) was used during both dissection and healing; no antibiotics were used. To optimize this protocol, neural plate of WT embryos was replaced with that of donor embryos previously injected with mRNA encoding myc-tagged GFP synthesized *in vitro* using Ambion’s mMessage mMachine kit (the template pCS2^+^-mt-GFP (BAMHI minus) was a gift from Michael Klymkowsky; Addgene plasmid # 15681) and we assessed the specificity of GFP expression in the spinal cord of resulting larvae.

#### *In vivo* Ca^2+^ imaging

Two- to four-cell-stage embryos were injected with 0.7-1 ng mRNA encoding the Ca^2+^ sensor GCaMP6s synthesized *in vitro* using Ambion’s mMessage mMachine kit (DNA template was subcloned into the pCS2^+^ vector as described in a previous study (Sequerra et al., 2018) from pGP-CMV-GCaMP6s, a gift from Douglas Kim, Addgene plasmid # 40753; Chen et al., 2013).

Embryos were grown to stage 39 at 23°C, then anesthetized for at least 10 min in 0.02% TMS and immobilized in a drop of 1% low-melting-temperature (26-30°C) agarose that was just above gelling temperature and placed at the bottom of a culture dish. Agarose was allowed to anneal, then MMR was added to the dish covering the agarose. Anesthetic was washed out for 10 min and larvae were time-lapse imaged for 15 min in 0.1X MMR/vehicle (0.1X MMR or 0.1% DMSO) with an acquisition rate of 0.5 Hz and a magnification of 25X using a fluorescent macroscope (Nikon AZ100). The temperature was set at 21°C during imaging using a Peltier thermoelectric cooler (QE-1HC and CL-100 from Warner Instruments).

After the first 15-min imaging period, larvae were anesthetized for 10 min in 0.02% TMS with or without a drug, then amputated at the distal edge of the imaged area (except for sham) corresponding to the midpoint of the tail (approximately 23 chevrons from the head), washed for another 10 min in saline with or without the drug, and the same area was re-imaged for 15-20 min. All cells within an 800 µm-long region of tail starting from the amputation site, excluding the fin, and showing any whole-cell, transient increases in fluorescence intensity were identified. The GCaMP6s fluorescence intensity of these cells was measured in both imaging periods using NIS-Elements software. Transients were detected as an increase in intensity greater than the standard deviation of the cell’s intensity when inactive multiplied by a number between 3.5 and 13 which was adjusted for each animal due to variable noise levels, but was consistent across cells and timepoints for individual larvae. Migratory cells were detected in some larvae but excluded from the analysis because previous research shows that these cells most likely were not neural or muscle progenitors. Muscle progenitors likely take much longer to migrate to the site of injury (Pipalia et al., 2016) and neural progenitors are derived from the spinal cord (Gargioli & Slack, 2004) which was not the source of the migratory cells observed in this study. In some larvae, the axial muscle of the tail contracted several times during the imaging period. To isolate the effect of the amputation on Ca^2+^ dynamics from the effect of muscle contractions, the time during the contraction through just after the signal returned to the initial intensity (typically 30-60 s) was eliminated from the analysis and the remaining data was normalized to the total imaging time. Statistical analysis was performed by calculating the change in the total number of transients in the first 15 min of imaging per larva before and after the amputation, then comparing this change across treatments. In active cells, the change in mean number of transients per cell per 15-min was also analyzed.

To image animals at 1 dpa, GCaMP6s-injected embryos were grown to stage 39 at 23°C, then anesthetized in 0.02% TMS and amputated. They were washed and grown at 21°C in 0.1X MMR for another 22-26 h, then embedded in agarose as described above and imaged for 10 min. The frequency of Ca^2+^ transients was calculated and compared among regions anterior and posterior to the amputation plane.

Grafted larvae with GCaMP6s expression only in the spinal cord were imaged at stage 42 to take advantage of the increased transparency of older larvae. Larvae were embedded in agarose as described above, time-lapse imaged for 20 min with a Sweptfield confocal microscope (Nikon) with an acquisition rate of 0.5 Hz and a 40X objective. Larvae were then incubated with 0.02% TMS and a drug(s) or vehicle-only for 10 min, amputated at the distal edge of the imaged area to remove the last third of the tail (excluding sham), washed for 10 min with drug(s) or vehicle, then re-imaged for 40 min in the presence of the drug(s) or vehicle. The time (median = 12 s) during axial muscle contractions was eliminated from the analysis and the remaining data was normalized to the total imaging time. Temperature was maintained at 21°C as described above. All regions of the spinal cord showing GCaMP6s expression were measured using IMARIS software and transients were detected in regions of interest with a diameter of 4.75 µm as changes in intensity greater than the 99^th^ percentile of all changes in intensity in these regions during the first imaging period for that larva.

#### Statistics

An initial analysis of the distributions of error for whole-mount immunostaining experiments lead us to conduct these experiments with 6 larvae per treatment group per experiment and at least 3 independent experiments per data set except for Pax7/pErk1/2 stainings which were performed with 3, and morpholino-injected that were performed with 9, larvae per treatment per replicate. N for *in vivo* imaging was not predetermined and each treatment was performed on 5-11 independent groups of larvae with no more than two larvae per treatment, and at least one control, per group. All data was tested for normality by Shapiro-Wilk and homogeneity of variance by Levene’s test prior to ANOVA. Two-way ANOVA were also tested for interactions. To test the effect of PD0325901 on regenerated notochord length, the mean for each level of treatment and replicate was calculated and analysis was performed by Friedman’s nonparametric analysis. The studies of the effect of PD0325901 on the number of Pax7+ cells in the regenerating tissues, the reduction in muscle regeneration by PD0325901, SHP099 and global morpholino, and of spinal cord regeneration by SHP099, the analysis of change in Ca^2+^ activity with global GCaMP6s acutely after amputation by distance from amputation, and the change in number of Ca^2+^ transients in grafted larvae all required a logarithmic transformation prior to ANOVA. The effect of spinal cord-only morpholino on muscle regeneration required a square-root transformation prior to ANOVA. The effect of PD0325901 and SHP099 on spinal cord length at 3 dpa required power transformations prior to ANOVA. These data are presented as back-transformed mean±95% confidence interval. All other averaged data are presented as mean±SEM between experiments unless otherwise noted. Post-hoc tests included Tukey, Tukey-Kramer, Dunnett’s, and Dunn’s means comparisons.

## FIGURE LEGENDS

**Figure 2-supplement 1.** Mek1/2 or Shp2 inhibition reduces active Erk1/2 in injured and regenerating tail. Stage-39 larvae were incubated immediately following amputation with either Mek1/2 (**A**; 3 µM PD0325901) or Shp2 (**B**; 50 µM SHP099) inhibitor or only vehicle (0.1 or 0.5% DMSO; control) for 2 or 24 h at 21°C, and then processed for whole-mount, pErk1/2 immunostaining. Images are representative maximum-intensity projections; scale bars are 100 µm. a: anterior, p: posterior.

**Figure 4-supplement 1**. Assessment of tissue regeneration in amputated larvae. Stage-39 larvae were amputated, allowed to regenerate for 1 to 5 days, then fixed and processed for whole-mount immunostaining for tissue-specific markers and imaged on a confocal microscope. Images are representative maximum-intensity projections. Arrowheads indicate the site of amputation. (**A**) Colocalization of pan-neuronal cell membrane marker (Xen1) and neural stem cells (Sox2) at 3 dpa. (**B**) Larvae were incubated following amputation with either vehicle-only (0.1% DMSO; control) or Mek1/2 inhibitor (10 μM PD98059) and organized muscle chevrons were counted under brightfield illumination 30 min post-amputation and again before fixation at 3 days post-amputation (dpa). Data are mean±SEM number of organized chevrons; N=5 experiments with n=7-9 larvae per treatment per experiment. (**C**) Muscle chevron organization through 5 dpa shown by 12/101 staining (mature skeletal muscle). The purple line delineates a single, organized muscle chevron. The region measured for regenerated skeletal muscle is outlined in red. (**D**) The amputation site (white arrowheads) was determined by notochord staining (Tor70) and morphology, and the length of spinal cord (red, dashed line; Sox2) and notochord were measured from this site. (**E**) Quantification of the regeneration timecourse for spinal cord (solid line; Sox2) and muscle (dashed line; 12/101) regeneration during the first 3 days at 21°C after amputation at stage 39. N≥3 experiments with n=3-6 larvae per treatment per experiment. Scale bars are 50 (**A, C**) or 100 (**D**) µm.

**Figure 4-supplement 2.** Erk1/2 activation is important for notochord regeneration. (**A**-**C**) Stage-39 larvae were incubated immediately following amputation with Mek1/2 inhibitor (0.1-3 µM PD0325901), Shp2 inhibitor (20 or 50 µM SHP099), or only vehicle (0.1 or 0.5% DMSO; control) for 3 days at 21°C and then fixed and processed for whole-mount immunostaining and imaged with a confocal microscope. (A)Representative maximum-intensity projections (MIPs). White arrowheads indicate amputation site; a: anterior, p: posterior; scale bar is 100 µm. Notochord was identified by Tor70 or MZ15 staining, or by morphology apparent in the background of Sox2 staining (red arrowhead indicates distal tip of notochord). (**B, C**) Data are mean+SEM length of the regenerated notochord measured in MIPs of the regenerating tail. N=3 experiments with n=4-6 larvae per treatment per experiment. Horizontal lines show the control mean and the gray areas show their errors. Statistical analysis was performed by Friedman’s nonparametric analysis on the experiment means, followed by Dunn’s multiple comparisons (B) or two-way ANOVA followed by Dunnett’s means comparison (**C**); *p<0.05, **p<0.001.

**Figure 4-supplement 3.** Mek1/2 inhibition with PD98059 reduces muscle regeneration. Stage-39 larvae were incubated immediately following amputation with Mek1/2 inhibitor (10 µM PD98059) or only vehicle (0.1% DMSO; control; 0 µM) and then grown for 5 days at 21°C. Whole larvae were processed for 12/101 (skeletal muscle) whole-mount immunostaining, and then imaged with a confocal microscope. (**A**) Representative maximum-intensity projections. Dashed lines delineate the border between stump and regenerated muscle; a: anterior, p: posterior; scale bar is 100 µm. (**B**) Data are mean+SEM sum of the area stained by 12/101 in the regenerating tail measured in each frame of the z-stack N=4 experiments with n=6 larvae per treatment per experiment. Triangles show means from independent experiments. Statistical analysis was performed by two-way ANOVA; **p<0.001.

**Figure 5-supplement 1.** Genetic knockdown of Mek1/2 and neural tissue grafting. (**A**-**D**) One-cell-stage embryos were injected with 3-11 pmol *map2k1.L*-targeted morpholino (*map2k1.L*-mo) without (**A, B**) or with (**C, D**) 4-13 pmol complementary, caging photo-morpholino (photo-mo), or with 9 pmol random control morpholino (control-mo; **A, B**), then grown to stage 22, 29, 32 or 39 (**A, B**), or 41 (**C, D**) and processed for Western blot assay of a single stage per experiment. (**A, C**) Representative full-length Western blots of whole-cell lysates pooled from 3 larvae per lane. (**C, D**) Injected animals and non-injected siblings (WT) were grown in the dark, then exposed (+UV) or not (-UV) to UV for 3 min 4-6 h prior to stage 39 to uncage the *map2k1.L*-mo, and then grown for another 24 h in the dark at 21°C. Results were corroborated in at least 4 independent experiments. (**B, D**) Band signal intensity was quantified using ImageJ. Data are mean±SEM percent difference in Mek1/2 signal between experimental and control lanes normalized to the loading control (Gapdh or alpha-tubulin). (**E**) The neural plate of stage 13-14 WT embryos was replaced with neural plate from siblings expressing GFP or GCaMP6s or containing *map2k1.L*-mo (schematic). Image shows a representative GFP-expressing neural tissue-grafted larva at stage 41.

**Figure 5-supplement 2.** Genetic inhibition of Erk1/2 signaling reduces notochord regeneration. One-cell-stage embryos were injected with 9-11 pmol *map2k1.L*-targeted morpholino (*map2k1.L*-mo) mixed with 11-13 pmol complementary, caging photo-morpholino. Animals were grown in the dark and then exposed to UV for 3 min 4-6 h prior to amputation at stage 39 to uncage the *map2k1.L*-mo, then allowed to regenerate for 3 days in the dark at 21°C, fixed and processed for whole-mount immunostaining, and imaged with a confocal microscope. (**A, C**) Images are representative maximum-intensity projections (MIPs) showing notochord identified by Tor70 staining. White arrowheads indicate amputation site; a: anterior, p: posterior; scale bar is 100 µm. (**A, B**) Controls are WT siblings exposed to UV and grown under the same conditions. (**C, D**) Larvae with *map2k1.L*-mo only in the spinal cord were generated by replacing the neural plate of stage 12.5-13 WT embryos with neural plate from their siblings with global *map2k1.L*-mo. Controls were generated by grafting neural plate from WT siblings onto WT embryos which were then exposed to UV and grown under the same conditions. Data are mean+SEM length of the regenerated notochord measured in MIPs of the regenerating tail. N=5-6 experiments with n=3-9 larvae per treatment per experiment. Scattered triangles show means of independent experiments. Statistical analyses were performed by two-way ANOVA; *p<0.05, **p<0.001.

**Video 1.** Stage-39 control larva globally-expressing GCaMP6s was imaged from 13 to 10 min prior to tail amputation with 25x magnification and 0.5 Hz. Playback at 40x and x-y resolution reduced to 10% of original.

**Video 2.** The same stage-39 control larva as in video 1 with global GCaMP6s expression was imaged from 10 to 13 min following tail amputation immediately anterior to the injury with 25x magnification and 0.5 Hz. Playback at 40x and x-y resolution reduced to 10% of original.

**Video 3.** Control larva globally-expressing GCaMP6s and imaged 1 day post-amputation (performed at stage 39) with 25x magnification and 0.5 Hz. Three min clip with playback at 40x and x-y resolution reduced to 10% of original.

**Video 4.** Stage-42 control larval expressing GCaMP6s only in the spinal cord was imaged from 13 to 10 min prior to amputation with 40x magnification and 0.5 Hz. Playback at 40x and x-y resolution reduced to 25% of original.

**Video 5.** The same stage-42 control larva as in video 4 expressing GCaMP6s only in the spinal cord was imaged from 10 to 13 min immediately anterior to the injury with 40x magnification and 0.5 Hz. Playback at 40x and x-y resolution reduced to 25% of original.

## ACKNOWLEDGEMENTS

We thank Neil Willits for his guidance with the statistical analysis of this data and Andrew M. Hamilton for his comments on the manuscript. This work was supported by NSF 1120796 and 1754340, NIH-NINDS R01NS073055, R01NS105886, R01NS113859 and Shriners Hospital for Children 86700-NCA grants to L.N.B.

