## Supplementary figures and images for "Injury-induced Erk1/2 signaling enhances Ca^2+^ activity and is necessary for regeneration of spinal cord and skeletal muscle"

### Figure 2-supplement 1

#### Figure 2-supplement 1

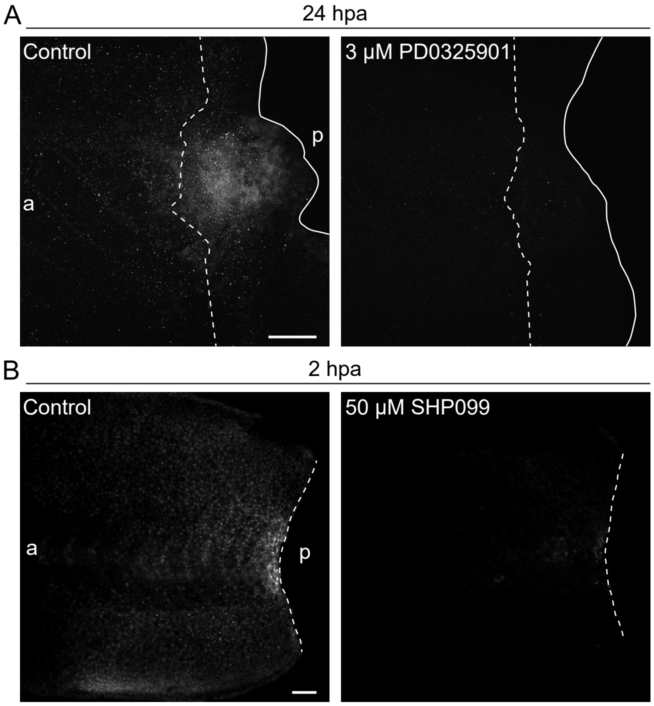

### Figure 4-supplement 1

#### Figure 4 - supplement 1

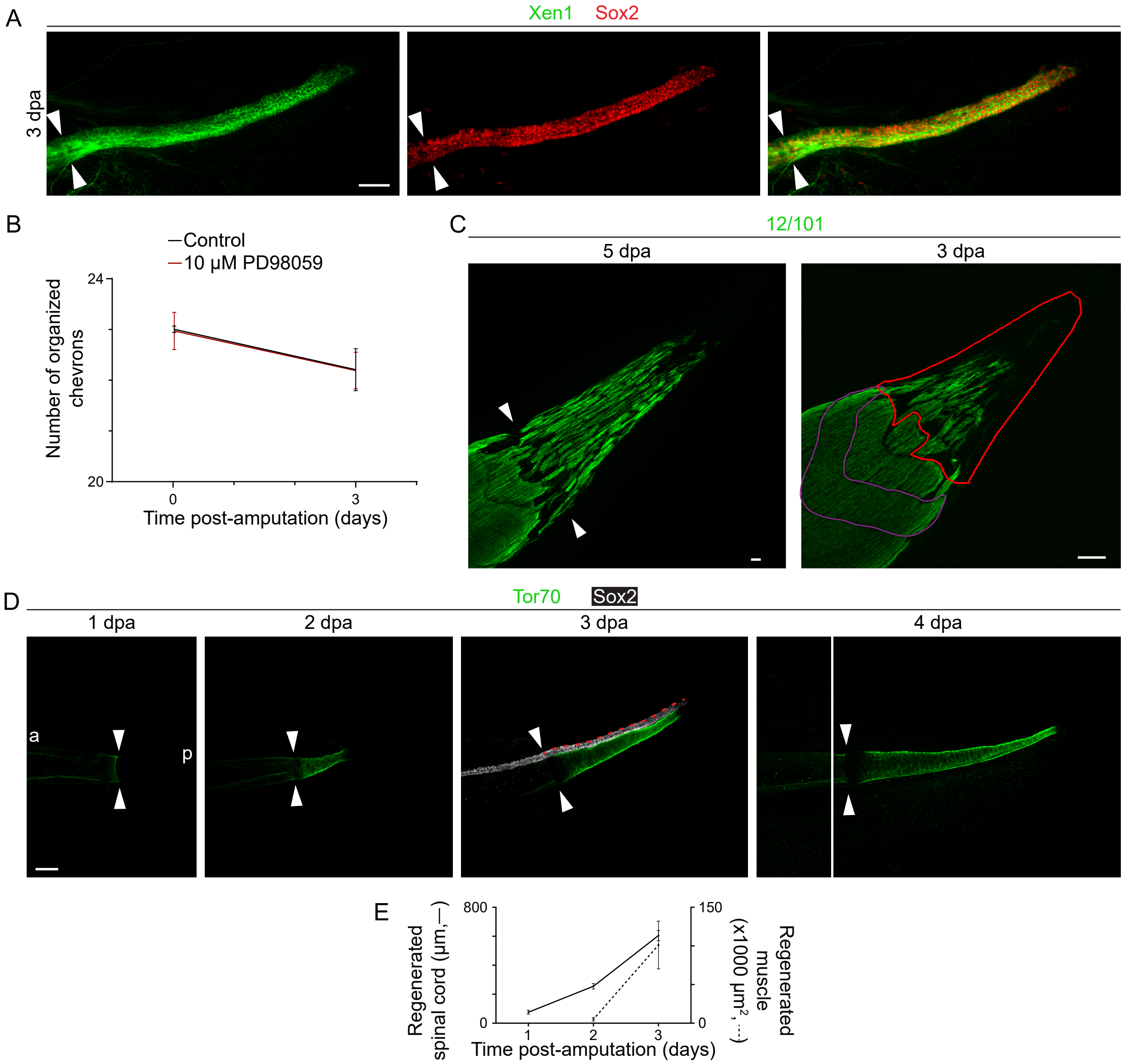

### Figure 4-supplement 2

#### Figure 4 - supplement 2

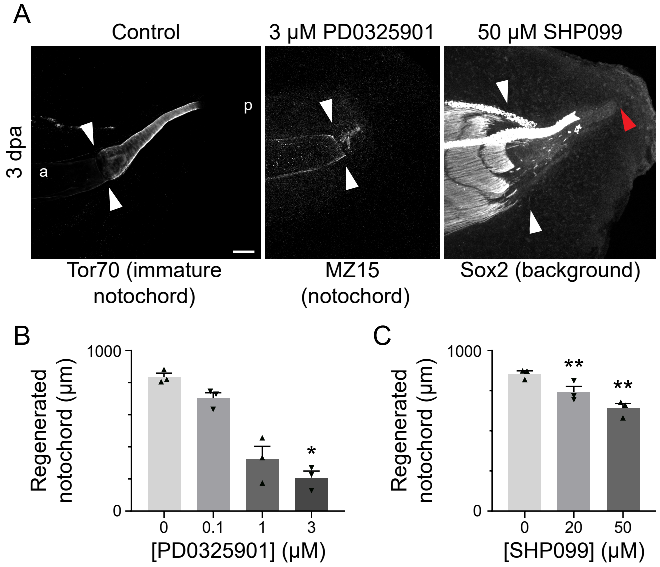

### Figure 4-supplement 3

#### Figure 4 - supplement 3

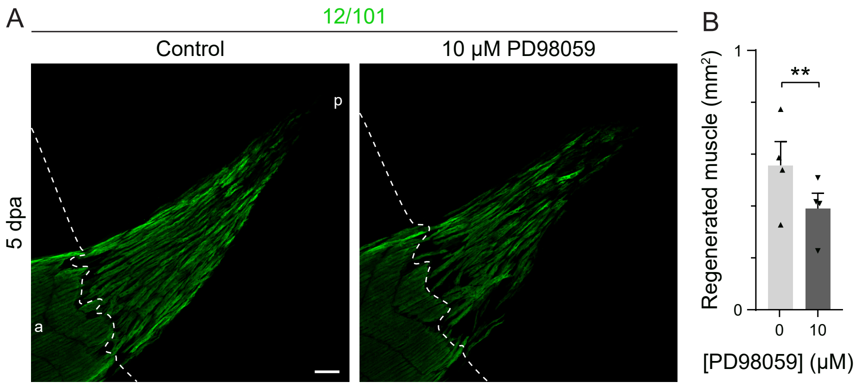

### Figure 5-supplement 1

#### Figure 5 - supplement 1

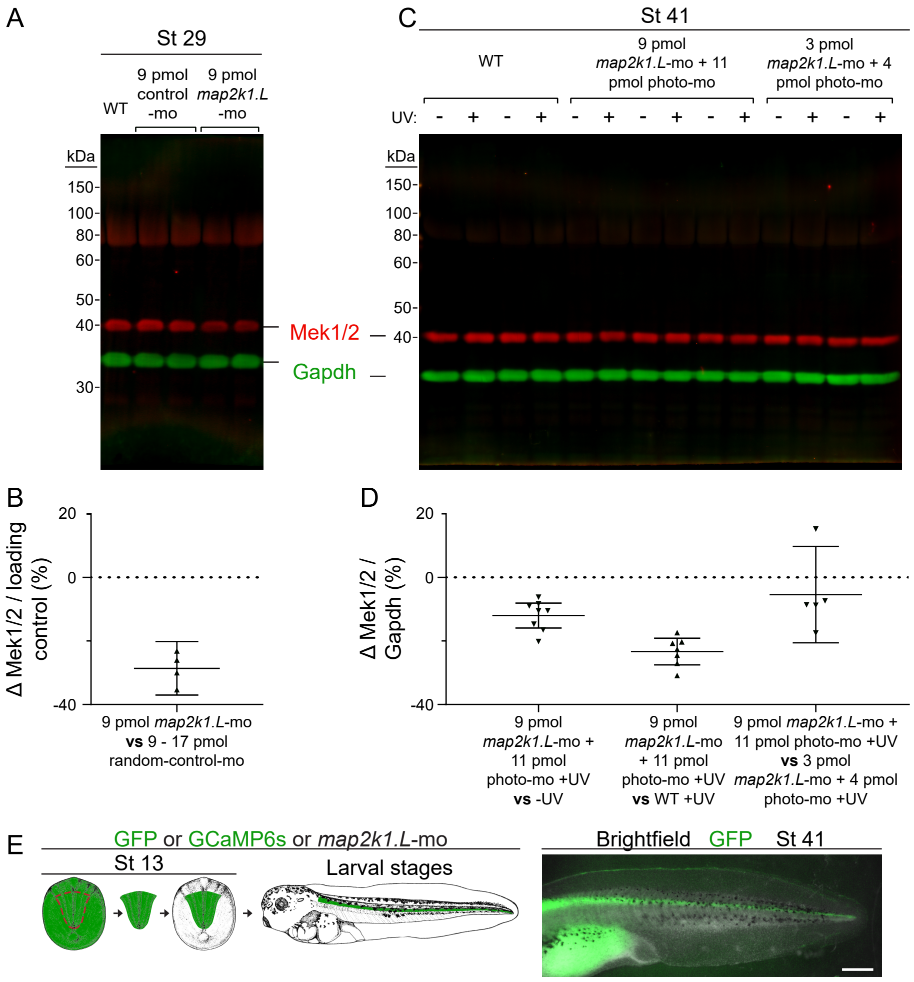

### Figure 5-supplement 2

#### Figure 5 - supplement 2

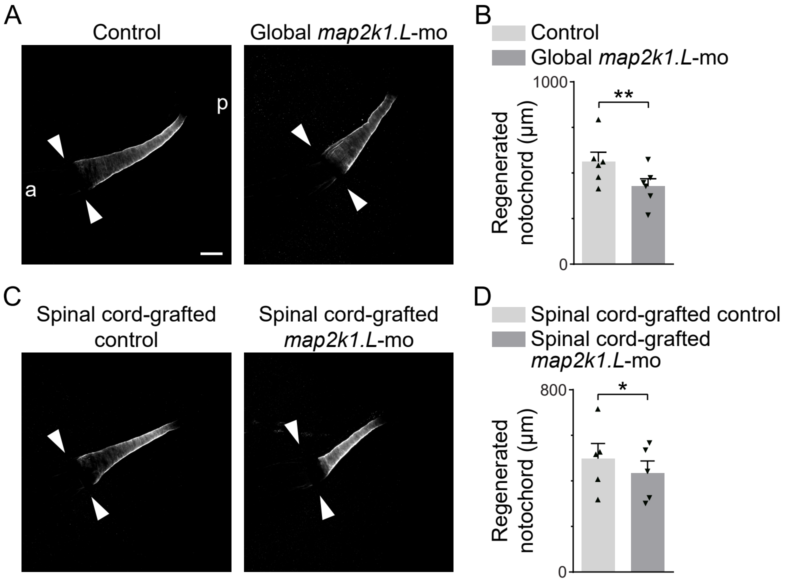
